# Acinar-ductal cell rearrangement drives pancreas branching morphogenesis in an IGF/PI3K-dependent manner

**DOI:** 10.1101/2023.01.26.525717

**Authors:** Jean-Francois Darrigrand, Anna Salowka, Francesca M. Spagnoli

## Abstract

During organ formation, progenitor cells need to acquire the diversity of cell identities found in the organ as well as organize themselves into distinct structural units. How these processes are coordinated, and how tissue architecture(s) are preserved despite the dramatic cell rearrangements occurring in developing organs remain unclear. Here, we identified cellular rearrangements between acinar and ductal progenitors as a mechanism to drive branching morphogenesis in the pancreas while preserving the integrity of the acinar-ductal functional unit.

Using *ex vivo* and *in vivo* mouse models, we found that pancreatic ductal cells form clefts by protruding and pulling on the acinar basement membrane, which lead to acini splitting. Newly formed acini remain connected to bifurcated branches generated by ductal cell rearrangement. IGF/PI3K pathway regulates this process by controlling ductal cell fluidity. If components of the pathway are genetically or chemically dysregulated, ductal cell fluidity prevents branching and affects pancreatic cell fates. Hence, our results explain how acinar multiplication and branch bifurcation are synchronized during pancreas organogenesis.

## INTRODUCTION

Numerous epithelial organs undergo branching morphogenesis during their development to build complex, arborized network and acquire specialized tissue-architectures. Examples include the lung, kidney and exocrine glands, such as the pancreas^1–3^. Understanding the establishment and maintenance of tissue architecture is a central question in developmental biology with a direct implication on organ physiology and disease. In fact, disruption of these same mechanisms in an embryo can result into developmental diseases^1,2^, while in adult life loss of tissue architecture may occur at early stages and across diverse human cancers^1,4^.

Live imaging approaches combined with organ cultures and genetic perturbations have provided invaluable insights into the molecular and cellular basis of branching. Several signaling pathways and cellular mechanisms have been identified as conserved regulators of branching morphogenesis in diverse organs, while important organ-specific differences have also arisen^1,4^. Recent studies have shown that each organ may employ a unique set of branching strategies to generate specialized shapes that need to be optimized for the organ’s physiological role^1,2,5,6^. Thus, the challenge is to decipher both ubiquitous and unique mechanisms.

The pancreas is a highly branched organ, wherein groups of acinar cells cluster around terminal ducts to form functional exocrine secretory units, and ductal cells line the tubular network of the pancreatic duct system^7,8^. In the mouse embryo, the formation of primary epithelial branches starts around E12.5, coinciding with the spatial segregation of pro-acinar and pro-ductal pancreatic progenitors^8–10^. Subsequently, a luminal plexus is remodelled into a tubular network in the center of the pancreatic epithelium, which is the site of endocrine differentiation, while the periphery displays ramifying branches with acinar cells placed at the ends of terminal ducts^11–15^. While early morphogenic events underlying primary branches initiation have been characterized to some extent^16–20^, there is no understanding of how acinar and ductal cells subsequently rearrange, when branches bifurcate to form a ramifying tree and acini multiply. Several fundamental questions remain unanswered about the coupling of differentiation and morphogenesis. It is indeed unclear the degree to which the morphogenetic potential of the tissue is determined by its differentiation state, as ductal-acinar units are already established while new branches are still being generated. Moreover, the molecular and cellular mechanisms ensuring branching morphogenesis while preventing detrimental ductal-acinar cell rearrangements are elusive.

Here, we addressed these questions by performing live-imaging and quantitative analyses of the cellular events underlying branching in the mouse embryonic pancreas. We found that ductal cells drive branching morphogenesis by a “protrude and pull” mechanism, which is IGF/PI3K-dependent. Our analyses of conditional *Igf1r* knockout mice and *ex vivo* pancreatic culture assays showed that ductal cells rearrangements are finely tuned by PI3K, enabling first clefting and splitting of the acini and then duct bifurcation. At the molecular level, PI3K activity depends on actomyosin contractility in ductal cells, highlighting an important role for tissue fluidity during branches formation in the pancreas. Together our findings explain how acinar multiplication and branch bifurcation are synchronized in the developing pancreas. They also strongly suggest that differentiated acini need to be established before bifurcating branches can be formed.

## RESULTS

### Ductal cells drive cleft-mediated branching morphogenesis in the pancreas

Here, we sought to elucidate the cellular mechanisms by which cells rearrange to support pancreas branching morphogenesis. To gain insight into these dynamic events, we first performed time-lapse imaging of ductal cells in mouse pancreatic explants collected at E12.5 (Figure 1A). *Ex vivo* pancreatic cultures closely recapitulate the *in vivo* morphogenesis processes, providing a simple platform for their observation (Figures S1A and S1B)^21,22^. We used the tamoxifen-inducible transgenic (Tg) *Krt19-* CreERT line^23^ in combination with the Tg(mTmG) fluorescent reporter line^24^ to mosaically label and track ductal cells in pancreatic explants. We observed that the Krt19-mG ductal cells localized in the centroacinar space often extend cellular protrusions between acinar cells and directed towards the basement membrane (BM) (Figure 1A; Video S1). Notably, our timelapse imaging showed that these ductal protrusions are dynamic, upon multiple cycles of contact formation and retraction they eventually pull the BM between neighboring acinar cells, initiating the formation of epithelial clefts (Figures 1B and S1C; Video S1).

**Figure 1.**
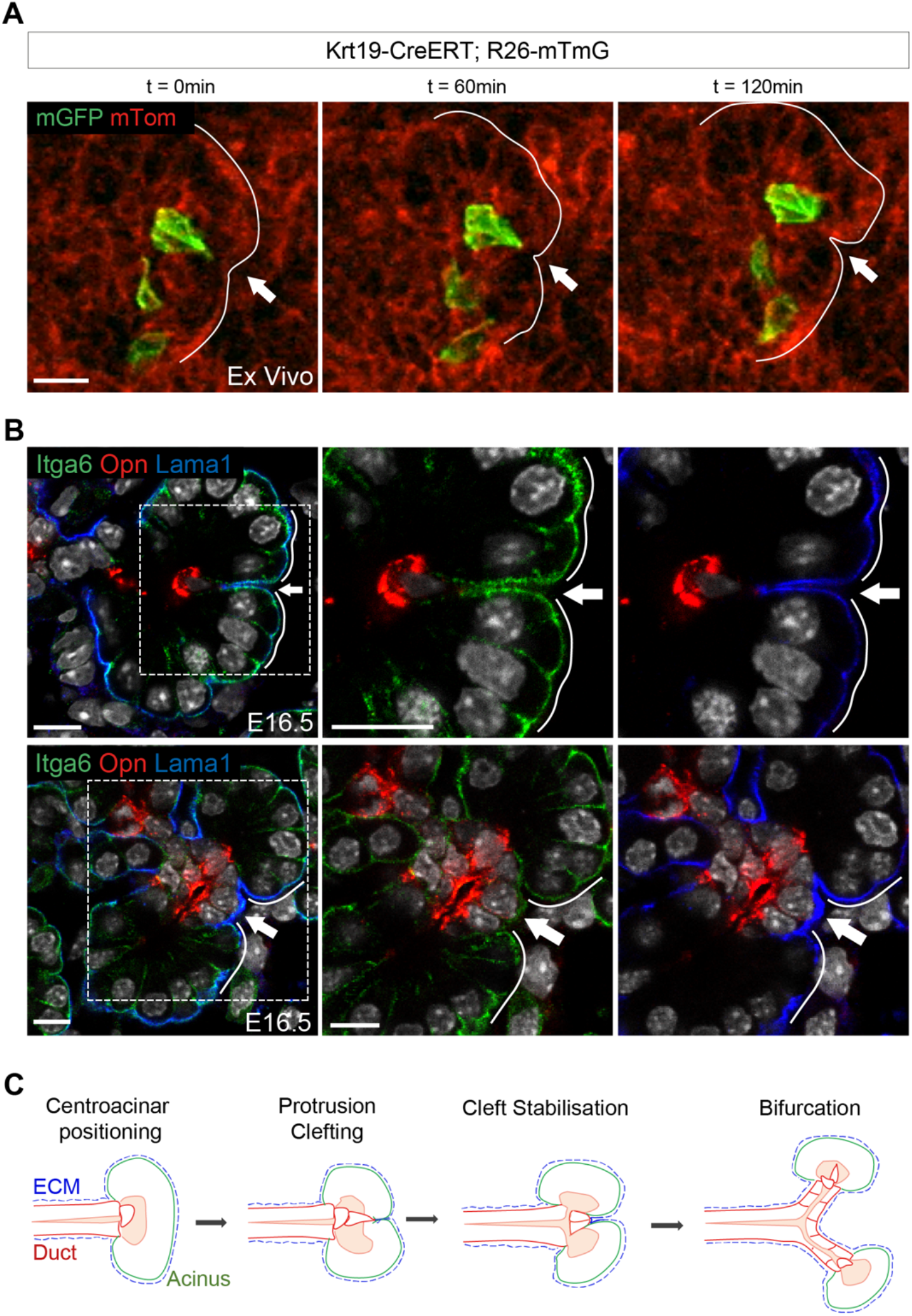
Pancreatic ductal cells protrusions promote cleft formation during branching morphogenesis. (A) Representative confocal time-lapse images of *Krt19-*Cre^ERT^;*R26*mTmG pancreatic explants collected at E12.5 and cultured for 48h in the presence of 4-hydroxytamoxifen (4-OHT). mGFP+ ductal cell (green) is shown protruding between acinar cells (red); arrow indicates a site of cleft formation. Time-lapse sequences of images from Video S1. Scale bar, 10 μm. (B) Representative confocal images of E16.5 pancreatic tissue immunostained for Integrin alpha6 (Itga6), Osteopontin (Opn) and Laminin alpha-1 (Lama1). Top panel, arrow indicates a cleft with a ductal cell (Opn+) underneath in contact with the BM (Lama1+); bottom panel, arrow indicates an enlarged cleft with three Opn+ ductal cells underneath. Boxed regions are shown at a higher magnification with split channels on the right. Scale bar, 10μm. (C) Schematic representation of ductal cells behavior driving the formation of clefts, the division of acini, and subsequently the bifurcation of the ducts.

To further assess whether the retraction of ductal protrusions might induce cleft formation, we performed immunofluorescence (IF) staining for markers of acinar (Integrin alpha6, Amylase, Cpa1), duct (Osteopontin, Sox9, Mucin, Cytokeratin 19) and BM (Laminin1) on pancreas tissue sections at E16.5, when many clefts are visible (Figures 1B and S1E). In all examined acinar clusters, we systematically found that below each cleft is positioned a ductal cell with protrusions anchored to the acinar BM that is invaginated between two acinar cells (Figure 1B). Following the formation of an initial cleft, the terminal ductal cells change shape, transitioning from a tear-drop shape into a cuboidal shape, and rearrange into a single-layer epithelium (Figures 1B and S1C). Concomitantly, neighboring ductal cells in contact with the protruding one appeared to enlarge the cleft by extending the interaction surface with the BM (Figures 1B and 1C; Video S2). These results suggest that a subset of ductal cells by a “protrude and pull” mechanism can drive ductal-mediated clefting and split the acini in two (Figures 1C and S1). BM digestion by collagenase treatment strongly reduced cleft numbers highlighting the importance of BM integrity for ductal-mediated clefting (Figure S1D). Moreover, our data showed that ductal-mediated clefting takes place from E14.5 onward (Figure S1F), which coincides with the period of secondary branching expansion^16,22^.

### PI3K regulates cleft formation and branching morphogenesis

The PI3K pathway is well known for regulating the ability of epithelial cells to protrude and rearrange during morphogenesis^25–28^. To investigate the potential influence of PI3K on cell rearrangement and ductal-mediated clefting during pancreatic morphogenesis, we treated explants collected at E12.5 either with the PI3K antagonist, LY294002, or the agonist, BpV(pic). After 24h of treatment, PI3K inhibition significantly increased the number of epithelial clefts, whereas PI3K overactivation decreased their number (Figures 2A and 2B). Notably, LY294002-induced clefting was accompanied by acini fragmentation, as evidenced by their significant decrease in size and increase in number (Figures 2A and 2C), while acinar cells number remained unaffected (Figure S2A). Conversely, PI3K overactivation caused an enlargement of the acini (Figures 2A and 2C). The morphological changes caused by PI3K dysregulation were not related to a change in cell proliferation, as the number of phospho-Histone 3 (PH3)^+^ cells was not significantly different in explants treated with LY294002 or BpV(pic) compared to non-treated controls (Figures S2B and S2C). However, high acinar cell death rate was detected in explants upon LY294002 treatment (Figures S2C and 2D).

**Figure 2.**
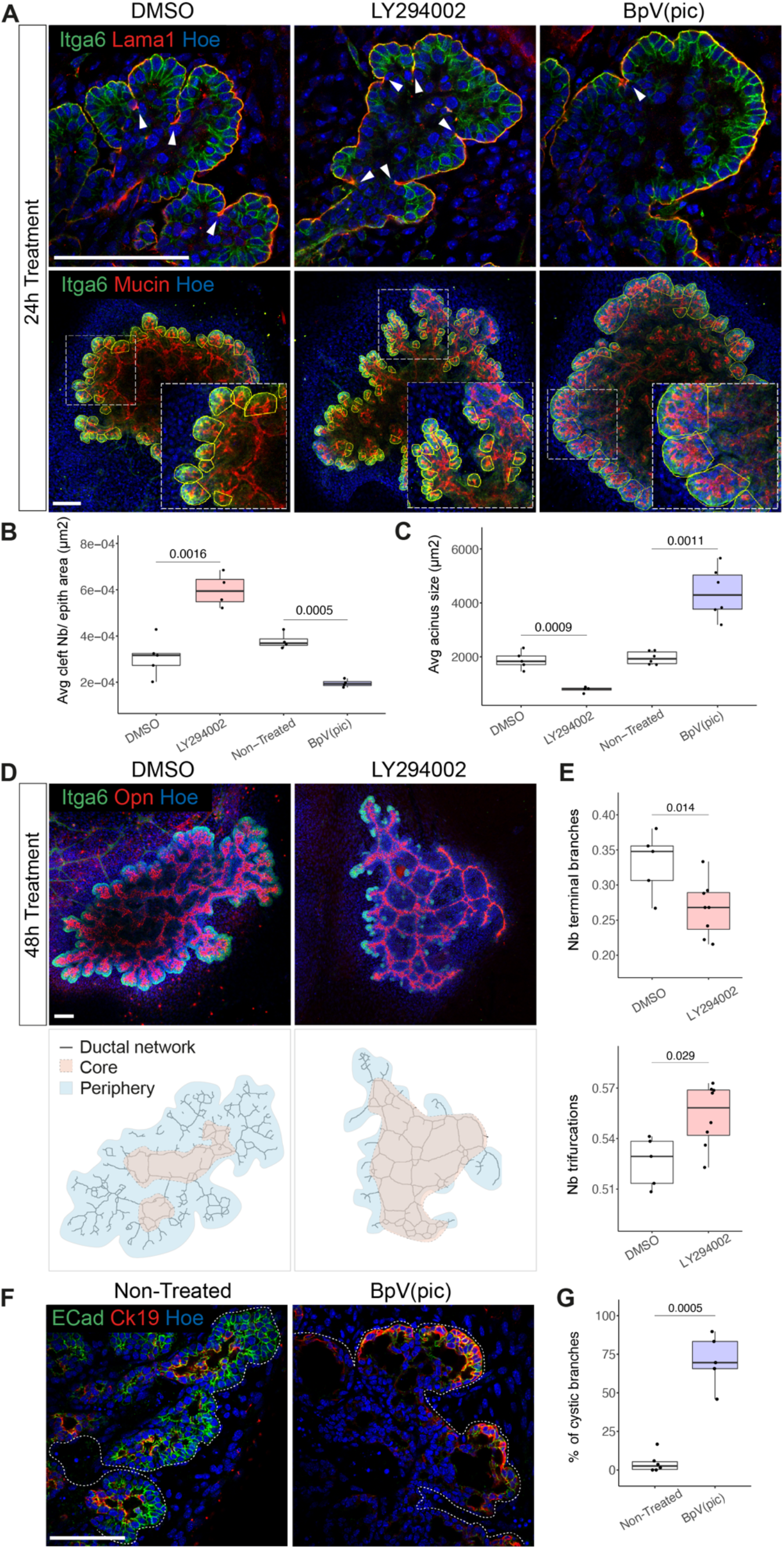
PI3K pathway regulates cellular rearrangements underlying branching morphogenesis. (A) Representative confocal images of pancreatic explants collected at E12.5, cultured for 24h in the presence of DMSO (Control), LY294002 or BpV(pic) and immunostained for indicated antibodies. Top: arrowheads indicate clefts. Bottom: acini are delineated with a yellow line and insets show higher magnifications of boxed regions. Scale bar, 100μm. (B) Quantification of clefts (in μm^2^) in pancreatic explants upon indicated treatments. The numbers (Nb) of clefts are shown relative to the area of the pancreatic epithelium (in μm^2^). n = 4-5 explants per treatment. Student’s *t*-tests. (C) Quantification of acini size (in μm^2^) in pancreatic explants upon indicated treatments. The numbers (Nb) of clefts are shown relative to the area of the pancreatic epithelium (in μm^2^). n = 4-6 explants per treatment. Student’s *t*-tests. (D) Top panel: Representative confocal images of pancreatic explants cultured for 48h with DMSO (Control) or LY294002 and immunostained for Itga6 and Opn. Scale bar, 100μm. Bottom panel: Binary ductal networks extracted from confocal images of the explants shown in top panel. Blue and beige overlays delineate the ductal network core and periphery regions, which are composed of interconnected and terminal ducts, respectively. (E) Quantification of terminal-ends (top) and trifurcations (bottom) in the ductal network of untreated and treated explants, expressed relative to the total number of branches. n= 5-8 explants per treatment. Student’s *t*-tests. (F) Representative confocal images of non-treated and treated pancreatic explants with BpV(pic) for 24h and immunostained for E-cadherin (Ecad) and Ck19. White dotted lines mark the pancreatic epithelium. Scale bar, 100μm. (G) Quantification of terminal cystic dilatations relative to the total number of branches. n= 5-6 explants per treatment. Student’s *t*-test.

During pancreas development, the ductal network grows following two distinct but concomitant morphogenetic programs. In the center (*a.k.a*. core), an expanding ductal mesh remodels to form a network of interconnected ducts, whereas in the periphery, terminal ducts grow and branch into a ramified network^11,29^. Upon long-term exposure to LY294002, we observed a major impact on the ductal network at the periphery, displaying a striking reduction of ramified branches as compared to the core area. To quantify this change in architecture, we extracted the ductal networks topologies of the explants^30^ (Figure 2D). Our quantitative analysis showed that PI3K inhibition strongly limits branching morphogenesis at the periphery with many ductal branches being devoid of acini, while central ducts remodeling in the core expanded (Figures 2D and 2E). Conversely, prolonged overactivation of PI3K transformed enlarged acini into cysts solely composed of ductal cells, as judged by the expression of Ck19 (Figures 2F and 2G). Together, our results indicate that prolonged PI3K inhibition induces first uncontrolled clefting and over-fragmentation of the acini, which subsequently lead to acinar loss and reduction of secondary and terminal branches.

### IGF1R acts upstream of PI3K in ductal cells to regulate branching

The PI3K pathway can be activated by a variety of ligands upon binding to Receptor Tyrosine Kinases (RTK) receptors at the cell membrane^27^. In order to identify which signals trigger PI3K activation in ductal cells, we analyzed the expression of RTK receptors in a publicly available scRNAseq dataset of pancreatic epithelial cells between E12.5 and E18.5^31^. We identified *Igf1r* as a promising candidate since it showed high levels of expression in the ductal cell cluster (Figure S3A). IF analysis confirmed the remarkably specific localization of IGF1R in ductal cells from E12.5 to E18.5 (Figures 3A and S3B). Conversely, the *Igf1* ligand showed abundant expression in acinar cells at the same stages, suggesting that acinar cells can send short-range signal to adjacent ductal cells in the centroacinar space (Figures 3B and S3A). 3D imaging by light-sheet fluorescence microscopy enabled to clearly visualize IGF1R membrane localization in ductal cells, decorating the protrusions and their intimate connection with the initiating clefts (Figure S3C; Video S3).

**Figure 3.**
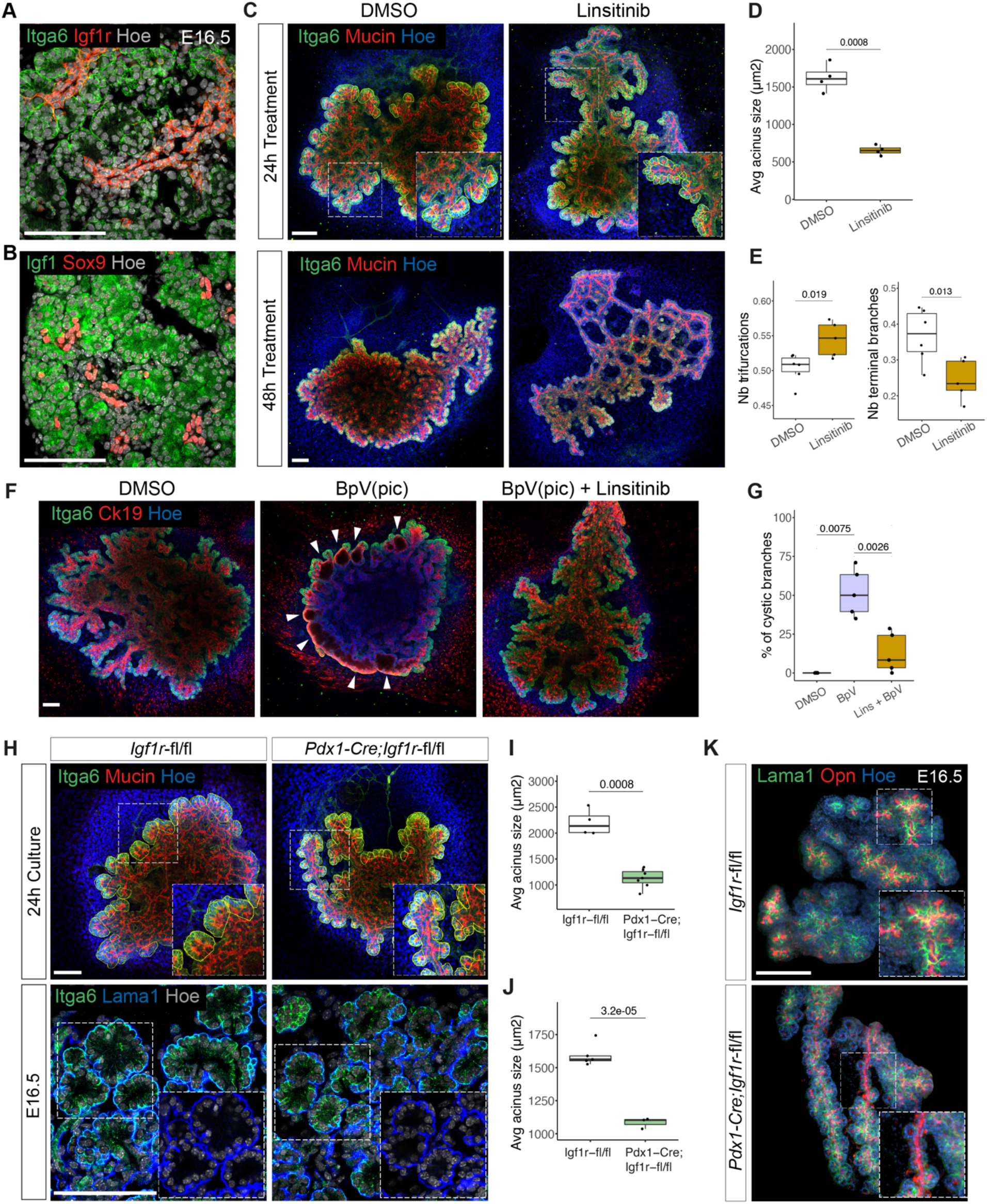
IGF signaling acts upstream of the PI3K pathway to regulate branching morphogenesis and acinar fragmentation. (A) Representative confocal images of pancreatic tissue at E16.5 immunostained for Itga6 and IGF1R.Scale bar, 100μm. (B) Representative confocal images of pancreatic tissue at E16.5 immunostained for IGF1 and Sox9. Scale bar, 100μm. (C) Representative confocal images of pancreatic explants collected at E12.5, treated for 24h (top panel) or 48h (lower panel) with DMSO (Control) or Linsitinib, and immunostained for Itga6 and Mucin. Top: insets show higher magnifications of boxed regions; yellow lines delineate acini. Scale bar, 100μm. (D) Quantification of acini size (in μm^2^) in pancreatic explants treated 24h with DMSO (Control) or Linsitinib. n= 4 explants per treatment. Student’s *t*-test. (E) Quantification of trifurcations (left) and terminal-ends (right) in the ductal network of explants treated 48h with DMSO (Control) or Linsitinib, expressed relative to the total number of branches. n= 5-7 explants per treatment. Student’s *t*-tests. (F) Representative confocal images of pancreatic explants collected at E12.5, treated for 48h with DMSO (Control), Linsitinib alone or in combination with BpV(pic), and immunostained for Itga6 and Ck19. Arrowheads indicate enlarged acini resembling cystic structures. Scale bar, 100μm. (G) Quantification of terminal cystic dilatations relative to the total number of branches. Scale bar, 100μm. n= 5 explants per treatment. Mann-Whitney *U*-test (DMSO vs BpV) and Student’s *t*-test (BpV *vs* Lins + BpV). (H) Representative confocal images of *Igf1r^flox/flox^* and *Pdx1-*Cre*;Igf1r^flox/flox^* pancreatic explants (top panel) and E16.5 pancreatic tissue (lower panel). Explants were immunostained for Itga6 and Mucin, pancreatic tissue for Itga6 and Lama1. Yellow lines delineate acini; insets show higher magnifications of boxed regions. Scale bar, 100μm. (I) Quantification of acini size (in μm^2^) in *Igf1r^flox/flox^* and *Pdx1-*Cre*;Igf1r^flox/flox^* pancreatic explants. n= 4-7 explants per genotype. Student’s *t*-tests. (J) Quantification of acini size (in μm^2^) in *Igf1r^flox/flox^* and *Pdx1-*Cre*;Igf1r^flox/flox^* E16.5 pancreatic tissue. n= 3-5 embryos per genotype. Student’s *t*-tests. (K) Representative light-sheet microscopy images of E16.5 *Igf1r^Flox/Flox^* and *Pdx1-* Cre*;Igf1r^flox/flox^* pancreata immunostained for Lama1 and Opn. Insets show higher magnifications of boxed regions highlighting unbranched ductal segments and partial loss of acini in the mutant pancreas. Scale bar, 1cm.

Next, to functionally probe whether IGF/IGF1R signaling is responsible for PI3K activation in ductal cells, we inhibited the pathway activity in pancreatic explants using linsitinib, an IGF1R antagonist. The blockade of IGF1R signaling led to a significant increase in acinar fragmentation (Figures 3C and 3D), which was followed by a reduced number of secondary branches at the periphery after 48h (Figures 3C and 3E). Thus, linsitinib treatment remarkably phenocopied PI3K inhibition, suggesting that IGF/IGF1R mediates the interaction between acinar and terminal ductal cells. To further test this hypothesis, we performed rescue experiments by blocking IGF1R activities in pancreatic explants exposed to the PI3K agonist, BpV(pic). Notably, the simultaneous treatment with linsitinib and BpV(pic) led to a significant decrease in the formation of cysts in comparison to explants treated with BpV(pic) alone (Figures 3F and 3G), indicating that IGF/IGF1R signals act upstream of PI3K activation in ductal cells. Additionally, we undertook a genetic approach to specifically knock-out *Igf1r*^32^ in the mouse pancreas epithelium (Figure S3D). *Igf1r* deletion in *Pdx1*-Cre*;Igf1r^flox/flox^* pancreatic explants caused a fragmentation of the acini *ex vivo*, like LY294002 and linsitinib treatments (Figures 3H and 3I), as well as *in vivo* in pancreatic tissue at E16.5 (Figures 3H and 3J). This acinar phenotype was coupled with defects in the bifurcation and ramification of the secondary ductal branches in the absence of severe pancreas growth defects (Figures 3K and S3E). On closer inspection, the mutant pancreata displayed long and unramified ductal segments that were sometimes devoid of acini (Figure 3K). Interestingly, double knock-out embryos for *Igf1r* and the highly homologous family member, *Insr*, showed stronger defects in pancreatic growth and branching morphogenesis^33^, suggesting functional compensation of the *Insr* in *Igf1r* mutant pancreas. Overall, our results underscore a role for the IGF signaling in regulating pancreas branching morphogenesis.

### PI3K finely tunes ductal cells fluidity to enable branching morphogenesis

To mechanistically understand how the IGF/PI3K pathway regulates ductal cells rearrangement to drive pancreas branching morphogenesis, we analyzed the organization and dynamics of the actomyosin cytoskeleton. We observed that PI3K inhibition significantly reduces phospho-myosin intensity in ductal cells, whereas PI3K overactivation increased it (Figures 4A and 4B). IF for F-actin showed similar results (Figures S4A and S4B). Besides the increase in phospho-myosin intensity on the apical side of ductal cells, PI3K overactivation triggered a partial relocalization of phospho-myosin to the basolateral side (Figure 4A). This was specific to ductal cells and was not observed in acinar cells (Figure S4C).

**Figure 4.**
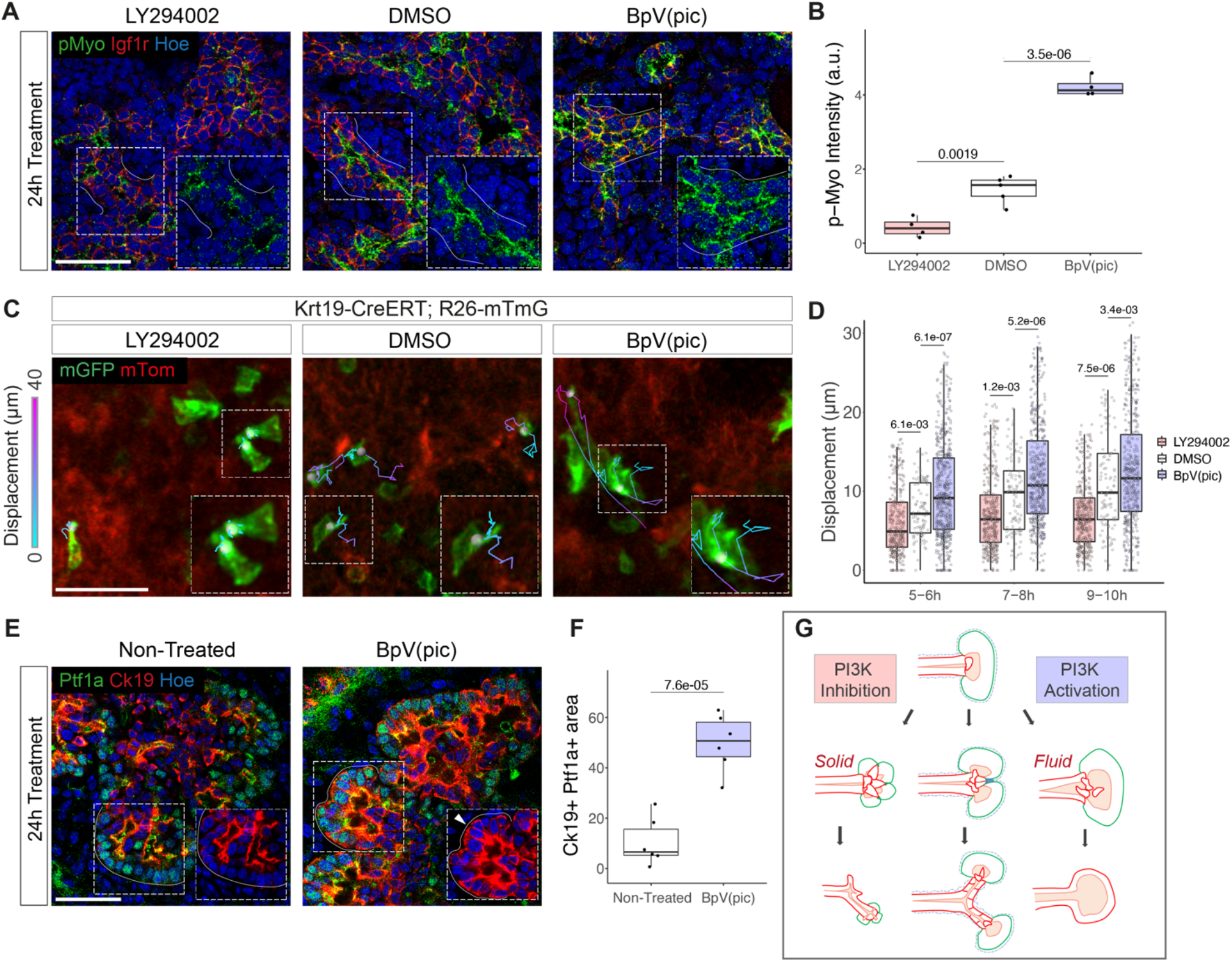
PI3K pathway regulates ductal cells fluidity in an actomyosin-dependent manner. (A) Representative confocal images of pancreatic explants sections collected at E12.5, treated 24h with DMSO (Control), LY294002 or BpV(pic) and immunostained for pMyo and Igf1r. Insets show higher magnifications of the pMyo channel in boxed regions. White lines delineate the basal side of ducts. Scale bar, 50μm. (B) Quantification of pMyo intensity at ductal cells membrane in pancreatic explants cryosections. n= 4-5 explants per treatment. Student’s *t*-tests. (C) Representative confocal time-lapse images of *Krt19-*Cre^ERT^*;R26mTmG* pancreatic explants collected at E12.5, cultured for 48h with 4-OHT and then treated with DMSO (Control), LY294002 or BpV(pic) and recorded for 12h. Representative tracks of mGFP+ ductal cells (green) taken from t=4h to t=8h frames of the time-lapse; color code corresponds to cell displacement (in μm). Scale bar, 50μm. (D) Quantification of ductal cells displacement (μm) in explants upon indicated treatments during a 2h-interval, ranging from 5h to 10h post-acquisition start. The displacement of hundreds of cells from n= 3 explants (per treatment) was analyzed. Student’s *t*-tests. (E) Representative confocal images of pancreatic explants non-treated or treated with BpV(pic) and immunostained for Ptf1a and Ck19. Insets show higher magnifications of the boxed regions. White lines delineate the basal surface of acinar cells; arrowhead indicates Ck19 localization in Ptf1a^+^ cells. Scale bar, 50μm. (F) Quantification of the area occupied by Ptf1a^+^Ck19^+^ cells in pancreatic explants upon indicated treatments relative to the total epithelial area (shown in %). n= 6 explants per treatment. Student’s *t*-test. (G) Schematic model of PI3K pathway regulation of pancreatic ductal cells fluidity, which in turn drives secondary branches formation.

Changes in actomyosin intensity and localization may translate into changes in tissue fluidity, as a result of the cells ability to move and exchange neighbors within an epithelium^34^. To assess whether PI3K might influence ductal cells fluidity, we followed ductal cell displacement using live imaging analysis in pancreatic explants upon exposure to LY294002 or BpV(pic). Cell tracking analysis showed that PI3K inhibition significantly impairs ductal cells displacement; many static cells were detected, displaying columnar shape and strong apical constriction (Figures 4C and 4D; Video S4). However, the LY294002 treatment did not affect protrusions formation, thereby ductal cells were still able to trigger initial clefts formation but not to subsequently rearrange and enlarge the clefts (Figures 4G and S4D). On the other hand, PI3K overactivation increased ductal cells displacement (Figures 4C and 4D; Video S4); many ductal cells showed loss of apical constriction and long-range migration, as in a fluid-like state. These results indicate that such uncontrolled ductal motility is responsible for the decreased ability of ductal cells to form clefts and for the disruption of acinar architecture (Figures 2A and 4G). In sum, ductal cells rearrangements appear to be finely tuned by PI3K, which enables cleft enlargement and duct bifurcation (Figure 4G).

Additionally, we found that these morphological changes were accompanied by altered acinar cell fate. After 24h of BpV(pic) treatment, a subset of Ptf1a^+^ acinar cells started to co-express ductal markers, such as Ck19, in the absence of acinar apoptosis (Figures 4E, 4F and S2C). Subsequently, after 48h, no acinar cells were detected in the explants (Figures S4E and S4F), suggesting that PI3K overactivation triggers an acinar-to-ductal transdifferentiation^35,36^. By contrast, we observed an increase in endocrinogenesis in LY294002-treated explants (Figures S4G and S4H). Overall, our results highlight that the PI3K-mediated regulation of ductal fluidity has a simultaneous impact on branching morphogenesis and pancreatic cell fate acquisition.

## DISCUSSION

Clefting at the branch tips in most branched organs is regulated by extrinsic mechanisms, including local physical constraints by external cell compression or ECM remodeling^6,37^. Our study offers a new perspective on pancreatic morphogenesis, whereby tissue fluidity would allow ductal cells to actively remodel the acinar compartment, pointing to an epithelial-intrinsic mechanism. We propose a two-step model of pancreatic branching morphogenesis, subsequent to the patterning of acinar and ductal domains in primary branches: 1) formation of clefts by ductal cell protrusions; 2) ductal cells rearrangement into secondary branches (Figures 1C and 4G). Overall, such a mechanism would enable the spatiotemporal synchronization between acini multiplication and ductal bifurcation, ensuring that acini remain all connected to the terminal end of ramifying ducts. Our findings also strongly suggest that during pancreas development the establishment of differentiated acini at the tip of primary branches is a prerequisite for branch bifurcation. Consistently, mutants with impaired acinar differentiation have strong branching defects, with ducts remaining as an interconnected mesh and lack of lateral branches^38^. Further studies are required to investigate if such a “protrude and pull” mechanism and a role for tissue-fluidity, described here in the pancreas, might extrapolate to other organs.

At the molecular level, we identify the IGF/PI3K pathway as a key regulator of cleft-mediated branching in the pancreas. We show that when PI3K is overactivated, ductal cells rearrange uncontrollably, which disrupts the epithelial architecture and results in the formation of cysts. The formation of cysts appeared to be linked with an acinar-to-ductal transdifferentiation. This result is in line with the consequences of PI3K overactivation in the adult pancreas that also triggers ductal cyst formation but via the expansion of centroacinar ductal cells instead of transdifferentiation^39^. This difference in cellular mechanisms could be due to differences between embryonic and adult tissues. Finally, the IGF system is intimately involved in the development and progression of pancreatic cancer^40^. If a similar IGF/IGFR acinar/duct embryonic niche might be exploited in metaplastic events underlying the initiation of pancreatic cancer is an open question, which deserves further investigation.

## Author contributions

J.F.D and F.M.S. conceived, coordinated the study, and wrote the manuscript with input from remaining authors. J.F.D. and A.S. performed and analyzed the experiments.

## Acknowledgements

We acknowledge the support of the European Union’s Horizon 2020 research and innovation programme Pan3DP FET Open [grant Number 800981]. JFD is currently supported by a fellowship from the NC3R [grant Reference NC/V002260/1].

## Declaration of interests

No competing interests declared.

## STAR METHODS

### RESOURCE AVAILABILITY

#### Lead Contact

Further information and requests for resources and reagents should be directed to and will be fulfilled by the lead contact, Francesca M. Spagnoli.

#### Materials Availability

This study did not generate new unique reagents.

#### Data and Code Availability

This study did not report any original code or RNA-sequencing data. Additional information on code used to complete this study is available from the lead contact upon request.

### EXPERIMENTAL MODELS

#### Animal work

All procedures relating to animal care and treatment conformed to the Institutional Animal Care and Research Advisory Committee and local authorities (PPL PP6073640, Home Office, UK). All mouse embryos were used without sex identification (mix sexes). The following mouse strains were used in this study: *Krt19-* Cre^ERT23^, *R26-*mTmG^24^, *Pdx1-*Cre^41^, *Igf1r*-flox^32^. All mice were bred on a C57BL/6 genetic background. Mice were housed in a specific pathogen-free facility in individually ventilated cages. Room temperature was maintained at 22 ± 1 °C with 30– 70% humidity and lighting followed a 12-h light/dark cycle. Food and water were provided ad libitum and none of the mice had been involved in previous procedures before the study. For timed mating, male and female mice were placed into a breeding cage overnight and plug check was performed daily. The presence of a vaginal plug in the morning was noted as E0.5.

## METHOD DETAILS

### Immunofluorescence staining

Embryos were collected at E12.5 to E18.5, the abdomen was dissected under a stereomicroscope in cold phosphate buffered saline (PBS) and fixed overnight at 4 °C in 4% paraformaldehyde (PFA). Fixed samples were washed in PBS and cryoprotected overnight in 20% sucrose. Tissues were embedded in OCT compound (Tissue-Tek, Sakura) and sectioned at 12μm thickness. For immunostaining, sections were incubated in TSA (Perkin Elmer) blocking buffer for 1h at room temperature (RT). If necessary, antigen retrieval was performed by boiling slides for 20 min in citrate buffer (Dako). Sections were incubated in primary antibody solution at the appropriate dilution overnight at 4°C. The following primary antibodies and dilutions were used: Amylase (1:400, Merck A8273), cleaved-Casp3 (1:300, Cell Signaling 9661), Collagen IV (1:200, Merck AB769), Cpa1(1:500, Biotechne AF2765), Cytokeratin19 (1:700, Abcam 133496), E-Cadherin (1:600, Merck U3254), Glucagon (1:500, Immunostar Inc. 20076), Igf1(1:300, Biotechne AF791), Igf1r (1:200, Biotechne AF305-NA), Insulin (1:2, Dako IR002), Integrin-α6 (1:400, Millipore MAB1378), Laminin-α1 (1:500, kindly provided by Prof. Sasaki, Oita University), Mucin (1:500, Thermo Scientific HM-1630-P1), Osteopontin (1:200, Biotechne AF808), phospho-Histone-H3 (1:300, Millipore 06-570), phospho-Myosin (1:200, Cell Signaling 3674), Ptf1a (1:200, kindly provided by Prof. Wright, Vanderbilt University), p120-Catenin (1:300, BD Biosciences 610134), Sox9 (1:500, Merck AB5535).

Sections were washed in PBS and incubated with a combination of Alexa fluor-conjugated secondary antibody (1:750, Invitrogen 11058, A11076, A21202, A21206, A21207, A21208, A31571, A31573; 1:750, Dianova 127-495-099) and counterstained with 250ng/ml of Hoechst 33342, for 1h at RT. When included, Phalloidin (1:400, Invitrogen A12379) was added to the secondary antibody solution. After washes in PBS, sections were mounted with SlowFade Gold Antifade Mountant (ThermoFisher Scientific). To perform multiple rounds of immunofluorescence labeling on the same tissue, antibodies were eluted by treating sections with 1% SDS solution (pH 2) for 1 h at 60°C. Slides were images on a Zeiss LSM 700 confocal microscope using 40×, 63× oil or 10× water immersion objectives.

### Explants culture

Dorsal pancreatic buds were microdissected from mouse embryos at E12.5 and cultured on glass-bottom dishes (Matek) coated with Fibronectin and filled with Basal Medium Eagle (BME) (10% FBS, 1% Glutamax, 1% Penicillin-Streptomycin, and 50 μg/ml Gentamicin)^21^. Explants were cultured for up to 72h in a tissue incubator (37°C, 5% CO2) and culture medium was changed daily. For explants treatments, BME culture medium was supplemented after 24h with the following small compound molecules: LY294002 (Biozol ST420-0005, final concentration 20 μM), BpV(pic) (Sigma SML0885, final concentration 2.5 μM), Linsitinib (Selleckchem S1091, final concentration 20 μM in Figure3 d-e and 40μM in Figure3 g-h) or Collagenase type IV (Sigma C5138, final concentration 20 μg/ml). Control explants were cultured with BME supplemented with the drug vehicle, DMSO, in the case of LY294002 and Linsitinib.

Following 24h or 48h of treatment, the explants were briefly washed with PBS, fixed for 20min at 4°C in 4% PFA and either processed for whole-mount immunofluorescence as previously described^21^ or equilibrated overnight in 20% sucrose and embedded in OCT for cryosectioning. Explants were imaged with a Zeiss LSM 700 confocal microscope.

### Time-lapse image acquisition

Dorsal pancreatic buds were microdissected from *Krt19-*Cre^ERT^;*R26*-mTmG embryos at E12.5 and cultured for 48h in BME supplemented with 4-Hydroxytamoxifen (4-OHT) (Sigma H6278, final concentration 1 μg/ml). Subsequently, 4-OHT was washed out and the explants were cultured in BME with or without the indicated small molecule compound for 1h before starting the acquisition. The microscope environmental chamber set-up was performed as previously described^21^. Time-lapse imaging of GFP+ ductal cells was performed overnight for 10-12h on a Zeiss LSM 700 confocal microscope, using a 10X water immersion objective and a viscous immersion fluid (Zeiss Immersol W). 8 μm interval Z-Stacks of 4 to 6 slices were taken every 10min.

### Light-sheet microscopy

Whole-mount immunofluorescence labeling of pancreata was performed as previously described^42^. For Igf1r labeling samples were treated with FLASH2 (80g/L Zwittergent, 250g/L Urea, in 200mM Borate) antigen retrieval solution prior to blocking^43^. Similar antibodies and dilutions were used as described above.

As described in Glorieux et al.^42^, CUBIC1 (25% wt/vol urea, 25% wt/vol N,N,N′,N′-tetrakis(2-hydroxypropyl) ethylenediamine, 15% wt/vol Triton X-100, in dH_2_O) and CUBIC2 (50% wt/vol sucrose, 25% wt/vol urea, 10% wt/vol 2,20,20′-nitrilotriethanol, 0.1% vol/vol Triton X-100, in dH_2_O) solutions were used for tissue clarification. Samples were mounted and imaged in a Zeiss Z1 light sheet microscope using 20X acquisition and 10X illumination lenses. 3D renderings were generated in Imaris software (Bitplane Oxford Instruments, version 9.5.1) using the “Crop 3D” tool on the area of interest and the “Surface” module.

### Image analysis

Structural features and immunostaining intensities were quantified on sections and explants using ImageJ^44^. The ductal networks topologies of explants were analyzed using the “Skeletonize (2D/3D)” and “Analyze Skeleton (2D/3D)” plugins (https://imagej.net/plugins/analyze-skeleton). Prior to the use of plugins 8-bit images of explants were processed as follows. The channel with Opn immunofluorescent labeling was split, a gaussian filter (sigma=2) was applied, the intensity was thresholded (Huang) to create a binary image, and small objects (below 300μm^2^ size) were automatically filtered out. From the data provided by the plugins, we extracted the number of End-points and Triple-points present in ductal networks and normalized them by the total number of branches. The End-points and Triple-points respectively represent a good approximation of the number of terminal branches and trifurcations in the ductal networks.

The membrane intensity of Phalloidin, E-Cadherin and phospho-Myosin were quantified by measuring the mean pixel intensity in manually drawn regions of interest (ROI) delineating ductal cells membrane. This value was normalized by the mean Hoechst intensity of each cell quantified and averaged per explant.

*Krt19*-GFP^+^ ductal cells were tracked in explants using Imaris. To quantify ductal cells displacement their surfaces were automatically generated using the “Surface” module on all focal planes (Z-Stacks of 4 to 6 slices with 8 μm interval). Filters were applied to discard multicellular objects and unspecific fluorescent speckles (“Number of voxels” superior to 100, “Area” comprised between 500 μm^2^ and 4300 μm^2^). Objects which were detected for less than 80 min or were detected discontinuously (gaps of more than 50 min present in their tracks) were filtered out. Any explant drift was corrected using the “Correct drift” tool on object tracks, which also corrected the image drift. The displacement of cells was measured using the “Detailed Statistics” tool and extracted as a .csv file. For generating the Video S4, we used the “Spots creation” module and manually edited the tracks.

### scRNAseq

The scRNA-seq data has been previously published in van Gurp et al.^31^. Uniform Manifold Approximation and Projection (UMAP) visualizations were imported from https://scdiscoveries.shinyapps.io/PancDev/.

### Quantification and Statistical Analysis

All statistical analyses were done using the R package rstatix, and statistical details related to tests performed and sample size are specified in the figure legends. For box-plots, the elements shown are the 25% (Q1, upper box boundary), 50% (median, black line within the boxes) and 75% (Q3, lower box boundary) quartiles, and the whiskers represent a maximum of 1.5X the interquartile range. p-values were calculated using two-tailed t-tests. Normality was assessed using the Shapiro-Wilk normality test. Statistical significance was calculated using Student’s *t*-tests for data displaying normal distributions and Mann-Whitney *U*-tests otherwise. In all experiments involving drug perturbations, experiments were repeated on explants collected from at least two embryonic litters.

---

**SUPPLEMENTAL INFORMATION**

**Figure S1.**
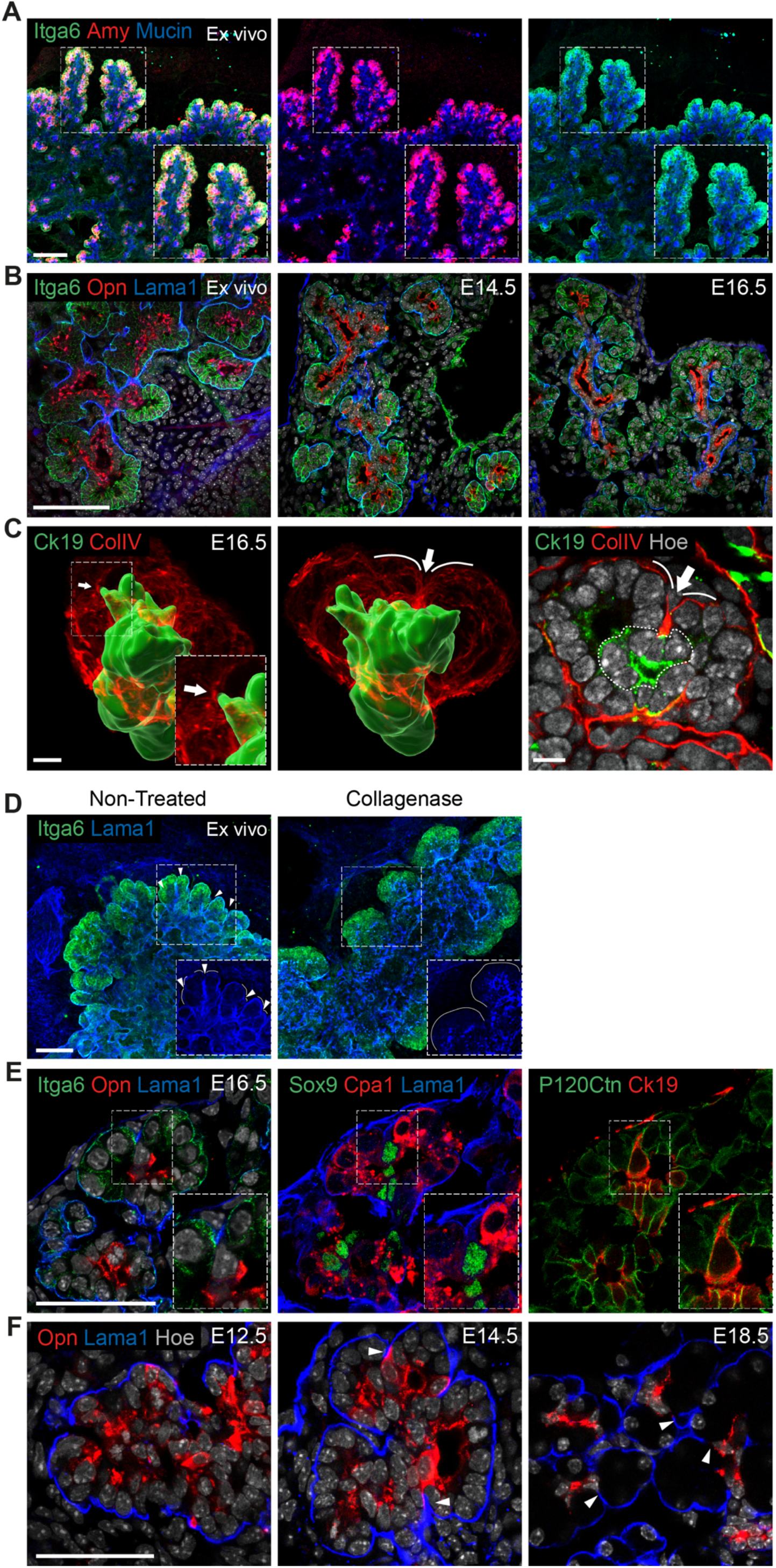
Characterization of pancreatic clefts and branches during embryonic development. (A) Representative confocal images of a pancreatic explant collected at E12.5, cultured for 48h and immunostained for Integrin alpha6 (Itga6), Amylase (Amy) and Mucin. Insets show higher magnifications of boxed regions. Different channel combinations are shown highlighting the colocalization between Itga6 and Amy in acinar cells. Scale bar, 100μm. (B) Representative confocal images of an *ex vivo* cultured pancreatic explant (left panel) and embryonic pancreatic tissue at E14.5 (middle panel) and E16.5 (right panel). All samples were immunostained for Itga6, Osteopontin (Opn) and Laminin alpha-1 (Lama1). *Ex vivo* cultures recapitulate the *in vivo* pancreatic tissue architecture, as previously reported^21,22^. Scale bar, 100μm. (C) Representative light-sheet images of E16.5 pancreatic tissue immunostained for Cytokeratin 19 (Ck19) and Collagen IV (ColIV). Left, Middle: 3D volumetric rendering showing Ck19^+^ ductal cells (green) and basement membrane (BM) (red). Arrows point at BM lining clefts in direct contact with a ductal cell below. Right: focal plane of the light-sheet Z-stack showing the 2D architecture of cleft. Scale bar, 50μm. (D) Representative confocal images of pancreatic explants non-treated or treated with Collagenase type IV and immunostained for Itga6 and Lama1. Arrowheads indicate cleft sites. Insets show higher magnifications of the boxed regions. Upon collagenase digestion, the BM appeared thinner and cleft number was reduced suggesting the importance of the BM during ductal-mediated clefting. Scale bar, 100μm. (E) Representative confocal images of multiplex immunostaining on E16.5 pancreatic tissue. The same tissue section underwent sequential immunostaining for different antibody combinations: Itga6 and Opn (left panel), Cpa1 and Sox9 (middle panel), p120Ctn and Ck19 (right panel). Insets show higher magnifications of the boxed region highlighting a protruding ductal cell positive for canonical ductal markers (Opn, Sox9, Ck19) positioned below a cleft. Scale bar, 50μm. (F) Representative confocal images of E12.5, E14.5 and E18.5 pancreatic tissue, immunostained for Opn and Lama1. Clefts (arrowheads) are visible at E14.5 and E18.5. Scale bar, 50μm.

**Figure S2.**
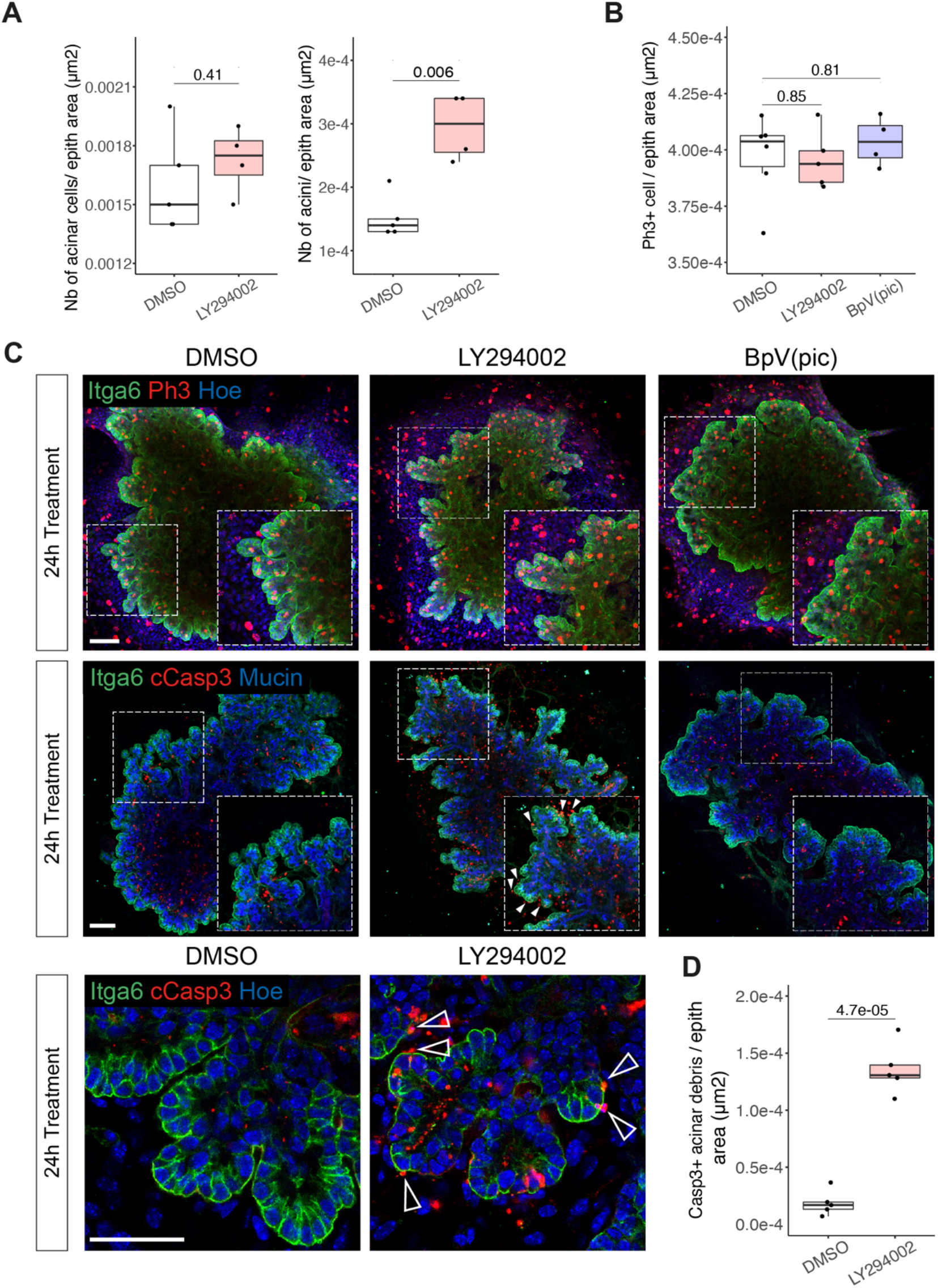
Impact of PI3K dysregulation on cell proliferation and apoptosis in pancreatic explants. (A) Quantification of number of acinar cells (left) and acini (right) in pancreatic explants treated with DMSO (Control) or LY294002 for 24h. Numbers are expressed relative to the epithelial area of the explants (in μm^2^). n= 4-5 explants per treatment. Student’s *t*-tests. (B) Quantification of Phospho-histone H3 (PH3)^+^ cells in pancreatic explants treated with DMSO (Control), LY294002 or BpV(pic) for 24h. Numbers are expressed relative to the epithelial area of the explants (in μm^2^). n= 4-6 explants per treatment. Student’s *t*-tests. (C) Representative confocal images of pancreatic explants collected at E12.5, exposed for 24h to indicated treatments and immunostained for Itga6 and PH3 (top) or Itga6 and cCasp3 (bottom). Insets show higher magnifications of boxed regions. cCasp3^+^ apoptotic debris (arrowheads) are detected next to the acini in LY294002-treated explants. Scale bar, 100μm. Bottom panel: zoomed images of pancreatic explants upon indicated treatments. Arrowheads indicate cCasp3^+^ debris extruded from acinar structures. Scale bar, 50μm. (D) Quantification of cCasp3^+^ apoptotic debris in contact with Itga6+ acinar cells in pancreatic explants upon indicated treatments. Values are expressed relative to the epithelial area of the explants (in μm^2^). n= 5 explants per treatment. Student’s *t*-test.

**Figure S3.**
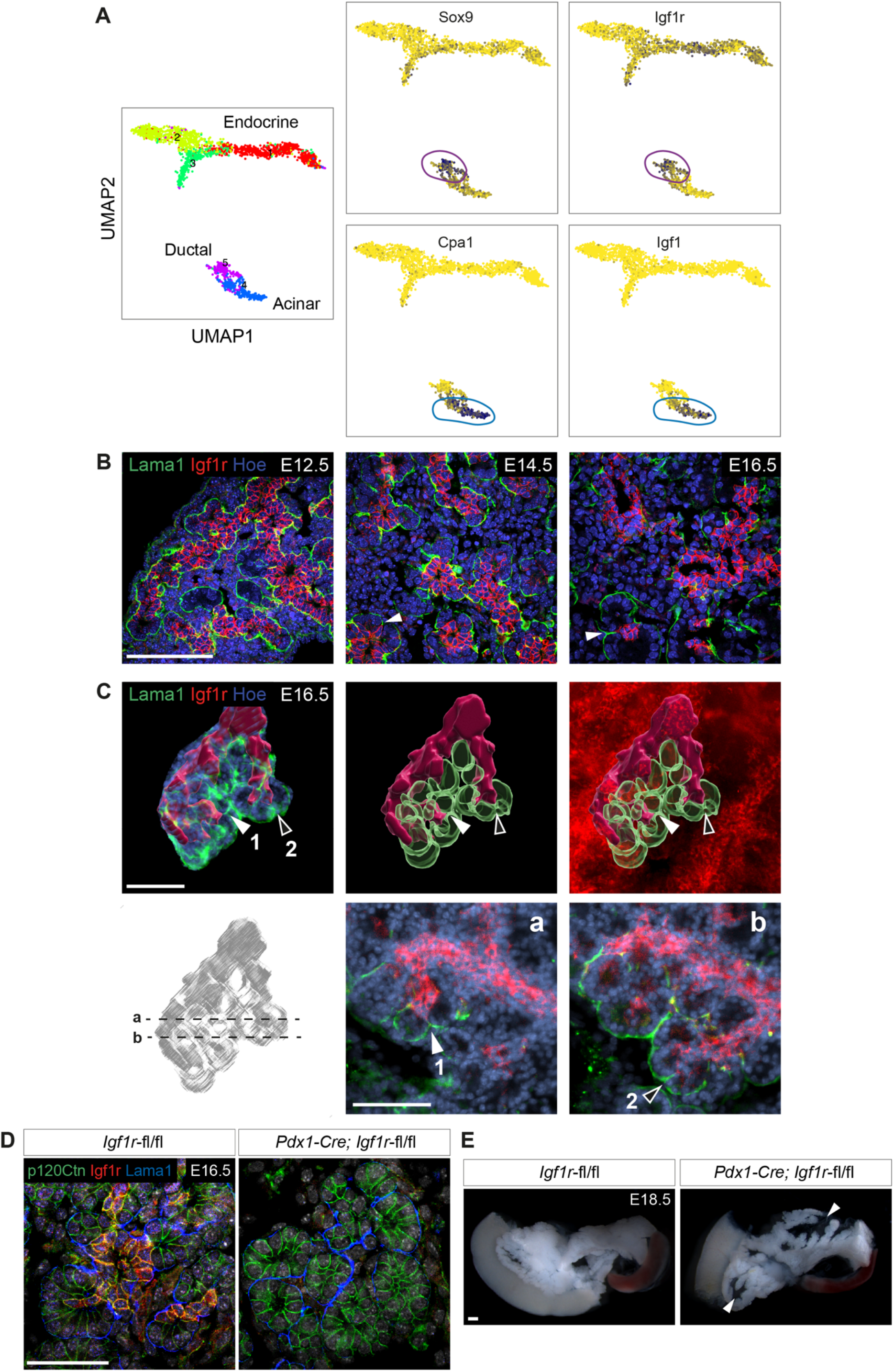
IGF1R specifically marks ductal cells in the developing pancreas. (A) Left panel: Uniform Manifold Approximation and Projection (UMAP) visualization representing pancreatic epithelial cells isolated from mouse embryos between stages E12.5 to E18.5 and color-coded based on their identity^31^. Right panels: Expression of genes marking the ductal (*Sox9* and *Igf1r*) and acinar (*Cpa1* and *Igf1*) clusters. (B) Representative confocal images of E12.5, E14.5 and E16.5 pancreatic tissue immunostained for IGF1R and Lama1. Clefts (arrowheads) are visible in E14.5 and E16.5 tissue. Scale bar, 100μm. (C) Representative light-sheet images of E16.5 pancreatic tissue immunostained for Lama1 and IGF1R. Top panel: 3D rendering showing IGF1R^+^ ductal structures (red isosurface) and Lama1^+^ BM surrounding the acinar structures (green isosurface). Arrowheads 1 and 2 point at clefts localized at the distal end of terminal branches. Bottom panel: focal planes of the light-sheet Z-stack showing the 2D architecture of cleft 1 (a) and cleft 2 (b). Scale bar, 100μm. (D) Representative confocal images of *Igf1r^flox/flox^* and *Pdx1*-Cre*; Igf1r^flox/flox^* pancreatic sections at E16.5 immunostained for p120Ctn, IGF1R and Lama1. *Pdx1-Cre;Igf1r^flox/flox^* mice showed a loss of *Igf1r* expression in the vast majority of pancreatic ductal cells. Scale bar, 50μm. (E) Representative stereomicroscope images of *Igf1r^flox/flox^* and *Pdx1-*Cre*;Igf1r^flox/flox^* pancreata at E18.5. Arrowheads indicate defects in long lateral branches in the mutant pancreas. Scale bar, 1cm.

**Figure S4.**
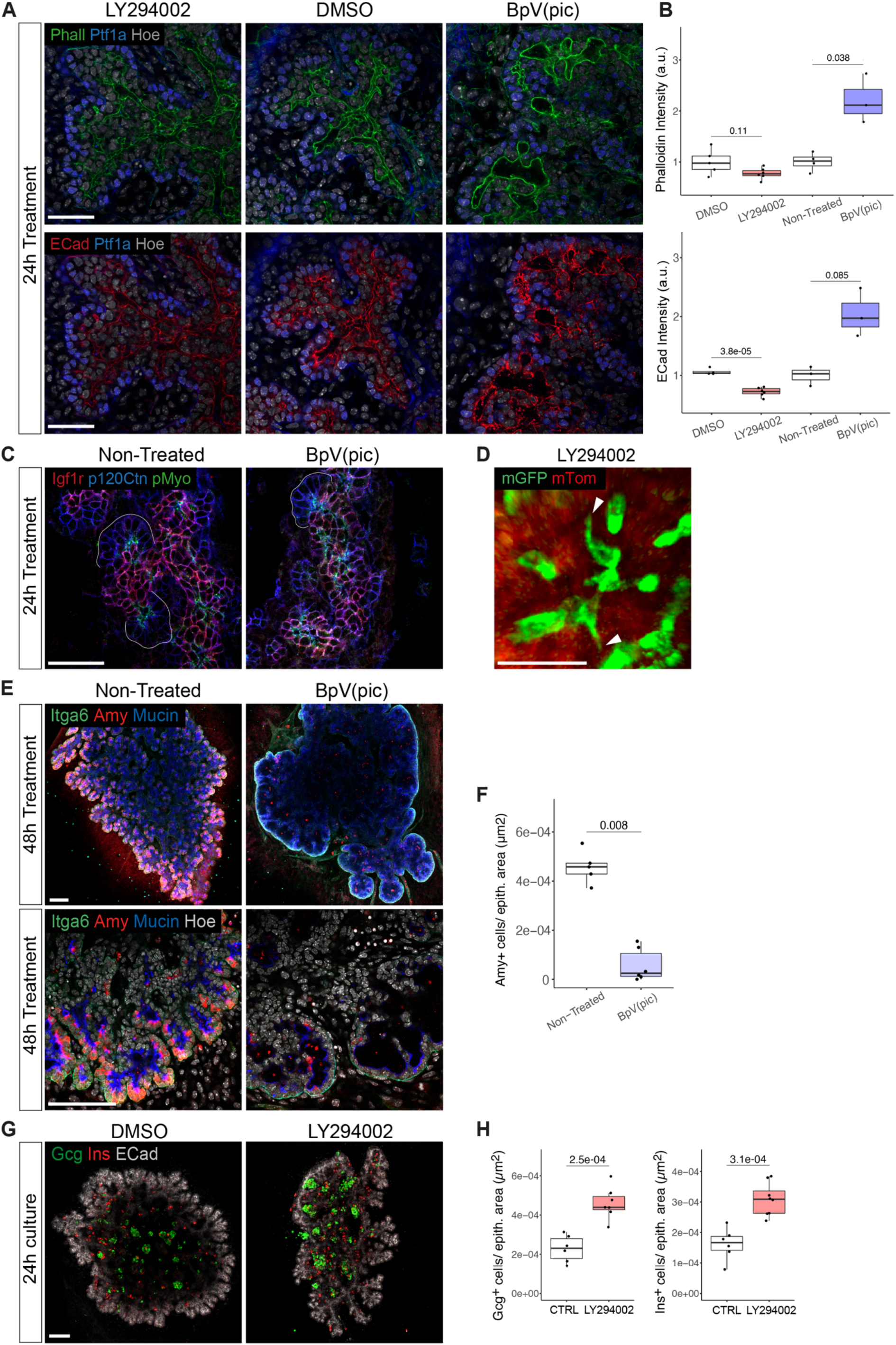
PI3K dysregulation affects actomyosin dynamics in ductal cells and pancreatic cell fate acquisition. (A) Representative confocal images of cryosections of pancreatic explants treated for 24h with DMSO (Control), LY294002 or BpV(pic) and immunostained for Phalloidin (Phall) (top panel), E-Cadherin (Ecad) (bottom panel) and Ptf1a. Scale bar, 50μm. (B) Quantification of Phalloidin (top panel) and E-Cadherin (bottom panel) intensities at the apical membrane of ductal cells in pancreatic explants upon indicated treatments. In this tissue-context, an increase in ductal fluidity did not coincide with a decrease in ductal cells adhesion. n= 3-6 explants per treatment. Student’s *t*-tests. (C) Representative confocal images of cryosections of pancreatic explants treated for 24h with BpV(pic) or non-treated (control) and immunostained for Igf1r, p120Ctn and pMyo. Scale bar, 50μm. (D) Representative confocal time-lapse images of *Krt19-*Cre;*R26*mTmG pancreatic explants treated for 24h with LY294002. Arrowheads indicates mGFP^+^ ductal cells (green) retaining the ability to form protrusions upon LY294002 exposure. Scale bar, 20μm. (E) Representative confocal images of pancreatic explants treated for 48h with BpV(pic) or non-treated (control) and immunostained for Itga6, Amy and Mucin. Bottom panel shows higher magnification of explants. Scale bar, 100μm. (F) Quantification of Amy^+^ cells in pancreatic explants upon indicated treatments. Numbers are expressed relative to the epithelial area of explants (in μm^2^). n= 5-6 explants per treatment. Mann-Whitney *U*-test. (G) Representative confocal images of pancreatic explants cultured for 24h with DMSO (Control) or LY294002 and immunostained for Glucagon (Gcg), Insulin (Ins) and ECad. Scale bar, 100μm. (H) Quantification of Gcg^+^ (left) and Ins^+^ (right) cells in pancreatic explants upon indicated treatments. Numbers are expressed relative to the epithelial area of explants (in μm^2^). CTRL, control. n= 6-8 explants per treatment. Student’s *t*-tests.

## LEGENDS OF SUPPLEMENTARY VIDEO

**Supplementary Video 1**

Confocal time-lapse video of *Krt19-Cre^ERT^; R26mTmG* pancreatic explants collected at E12.5 and imaged after 48h of 4-Hydroxytamoxifen (4OHT) treatment. mGFP+ ductal cells (green) are shown extending protrusions in between acinar cells (red). The first protrusion (first arrow) does not lead to epithelial clefting, while the second one does (second arrow). White line delineates the basal side of acinar cells, where the basement membrane (BM) lies.

**Supplementary Video 2**

Light-sheet fluorescent microscopy images of ductal branch overlying acini from E16.5 pancreas immunostained for ColIV (red) and Ck19 (green). Part 1: 3D rendering of the tissue, Ck19 isosurfaced in green. First arrow indicates the initiation of a cleft; second arrow points at an enlarged cleft, which separates two acini. Part 2: Z-stack visualization of the area with the enlarged cleft (third arrow).

**Supplementary Video 3**

3D rendering of E16.5 pancreatic tissue immunostained for Igf1r (red) and Lama1 (green) and imaged by light-sheet fluorescent microscopy. Selected region of interest displaying the organization of the Igf1r^+^ ducts (red isosurface) with respect to the acini (Lama1^+^ BM, green isosurface). Nuclei were stained with Hoechst (blue).

**Supplementary Video 4**

Part 1: Confocal time-lapse video of *Krt19-Cre^ERT^;R26mTmG* pancreatic explants cultured for 48h with 4-OHT, imaged for 12h after DMSO exposure. Tracks of mGFP^+^ ductal cells (green) from t=4h to t=8h time-lapse interval are displayed. Tracks are color-coded to illustrate cell displacement (in mm). In red, mTom^+^ unrecombined epithelial cells.

Part 2: Confocal time-lapse video of *Krt19-Cre^ERT^;R26mTmG* pancreatic explants cultured for 48h with 4-OHT, imaged for 12h after LY294002 exposure. Tracks of mGFP^+^ ductal cells (green) from t=4h to t=8h time-lapse interval are displayed. Tracks are color-coded to illustrate cell displacement (in mm). In red, mTom^+^ unrecombined epithelial cells.

Part 3: Confocal time-lapse video of *Krt19-Cre^ERT^;R26mTmG* pancreatic explants cultured for 48h with 4-OHT, imaged for 12h after BpV(pic) exposure. Tracks of mGFP+ ductal cells (green) from t=4h to t=8h time-lapse interval are displayed. Tracks are color-coded to illustrate cell displacement (in mm). In red, mTom^+^ unrecombined epithelial cells.

## Notes

### Competing Interest Statement

The authors have declared no competing interest.

